# Metabolization of α-D-carba-glucosamine in vivo generates antimetabolites of cell wall precursors

**DOI:** 10.1101/2023.02.08.527593

**Authors:** Milena Mund, Simon Friz, Anna Esser, Daniel Matzner, Alexander Babczyk, Dirk Menche, Heike Brötz-Oesterhelt, Christoph Mayer, Günter Mayer

## Abstract

α-D-Carba-glucosamine (CGlcN) is a carbocyclic analog of α-D-glucosamine that inhibits growth of *Bacillus subtilis* and *Staphylococcus aureus*. CGlcN is internalized and concomitantly phosphorylated via the phosphotransferase system yielding α-D-carba-glucosamine-6-phosphate (CGlcN6P), which interferes with expression of the glutamine-fructose-6-phosphate amidotransferase (GlmS; glucosamine synthase) by activating the *glmS* riboswitch. Herein, we report that CGlcN6P is efficiently metabolized to carbasugar nucleotides along the peptidoglycan biosynthetic route. Mass spectrometric analysis confirmed the occurrence of carbocyclic peptidoglycan nucleotides UDP-carba-D-*N*-acetyl-glucosamine (UDP-CGlcNAc) and UDP-carba-D-*N*-acetylmuramic acid-pentapeptide (UDP-CMurNAc-5P) in the presence of CGlcN and revealed accumulation of these carba-metabolites upon antibiotic treatment interfering with biosynthetic enzyme functions. Thus, carbocyclic carbohydrates and nucleotide analogs are generated by the promiscuous bacterial cell wall biosynthetic enzymes and act as antimetabolites, causing bacterial growth inhibition by interference with cell wall synthesis. Our findings reveal CGlcN not only as putative antibiotic molecule with previously unknown antimetabolite mode of action, but also as tool to study the bacterial cell wall metabolism, e.g., in synergy with other antibiotics.

---

Carbasugars are alicyclic carbohydrate analogs that contain methylene in place of the ring oxygen. They structurally resemble the parental sugars, but are unable to perform the acetal chemistry of their carbohydrate analogues. This increases stability and makes carbasugars useful probes as competitive inhibitors of enzymes such as the glycosidases and glycosyltransferase.^1^ Recently, an efficient, high-yielding synthesis of α-D-carba-glucosamine (CGlcN) was reported that allowed the biological evaluations of its mode of action as an glucosamine analog.^2–5^ In *B. subtilis*, CGlcN is taken up and concomitantly phosphorylated, like the natural metabolite, via the glucosamine-specific PTS system GamP, gaining intracellularly CGlcN-6-phosphate (CGlcN6P).^3^

Due to impaired acetal chemistry CGlcN6P cannot be deamidated via the glucosamine deamidase (NagB) such as the carbohydrate counterpart glucosamine-6-phosphate (GlcN6P).

However, due to a feedback mechanism, it regulates the expression of GlmS, an essential enzyme that catalyzes GlcN6P synthesis.^3^ This regulation is mediated by the *glmS* ribozyme, located in the 5’UTR of the *glmS* mRNA. Self-cleavage of the *glmS* ribozyme is activated by GlcN6P binding, which ultimately leads to a hydrolysis of the mRNA.^6–9^ CGlcN6P also is able to activate the *glmS* ribozyme in vitro and in vivo, resulting in a loss of *glmS* mRNA and bacterial growth inhibition.^3^

We previously demonstrated that (5a*R*)-Fluoro-α-D-carba-glucosamine-6-phosphate (F-CGlcN6P), another synthetic carba-mimic of GlcN6P, also activates the *glmS* riboswitch *in vitro*.^10^ This activation, however, was found to be ~32fold less efficient when compared to CGlcN6P (EC_50_-values are 192 μM vs. 6 μM). Treating *B. subtilis* with (5a*R*)-Fluoro-α-D-carba-glucosamine (F-CGlcN), a precursor of F-CGlcN6P, revealed inhibition of bacterial growth and we determined a minimal inhibitory concentration (MIC) of ~40μg/mL (**Fig. 1a,b**). Interestingly, this value is in the same range as the one found for CGlcN (32 μg/mL) although the activation of the *glmS* riboswitch *in vitro* was much less effective with F-CGlcN6P. We further demonstrate that antibacterial activity of F-CGlcN depends on uptake by GamP and induces cell envelope stress (**Fig. 1c,d**), measured by experiments using reporter genes under control of the global stress signals promoters.^11–14^ These two features are also observed from treating *B. subtilis* with CGlcN.^3^ These contradicting data suggest a secondary antibacterial effect of CGlcN (and F-CGlcN) which might be independent of *glmS* riboswitch activation.

**Figure 1.**
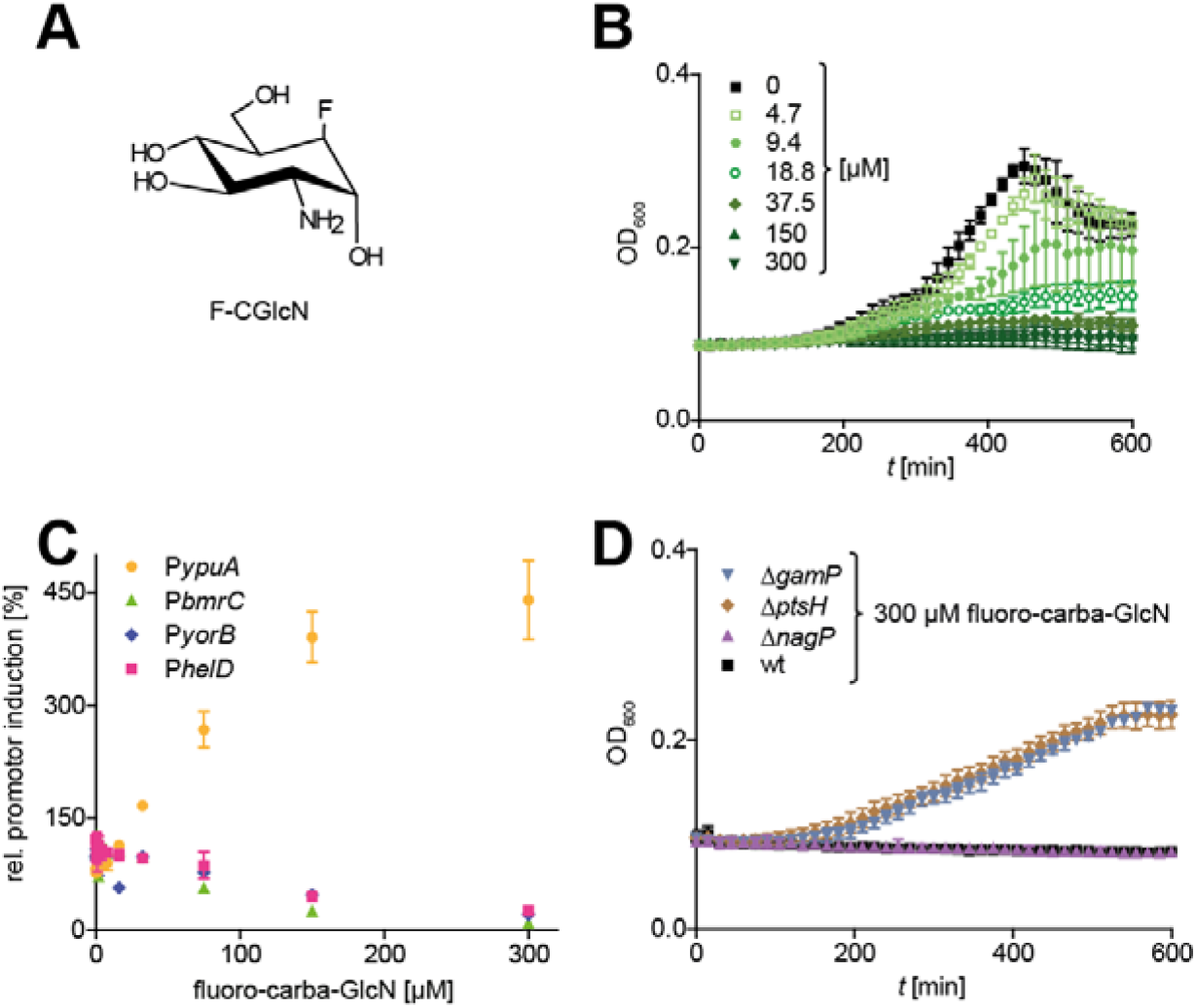
Impact of (5aR)-Fluoro-α-D-carba-glucosamine (F-CGlcN) on growth of wild-type *B. subtilis* and on mutant strains as well as stress response in reporter strains. *(A)* Chemical structure of Structure of F-CGlcN. *(B)* Growth curves of *B. subtilis* 168 in the presence of F-CGlcN (0–300 μM) were obtained from optical density at 600 nm. *(C)* Cell envelope stress response after F-CGlcN treatment shown by firefly luciferase expression under the control of stress-inducible promotor ypuA. No induction of bmrC promotor (protein synthesis inhibition), held promotor (RNA biosynthesis inhibition) and yorB promotor (DNA synthesis inhibition/damage) were observed. *(D)* Impact of deletions of *gamP* (blue; encoding the glucosamine PTS), *ptsH* (brown; encoding the general PTS kinase HPr) and *nagP* (purple; encoding the *N*-acetylglucosamine PTS) on the antibacterial activity of F-CGlcN compared to wt *B. subtilis* 168 strain.^15–17^ Growth curves were measured in the presence of 300 μM F-CGlcN as described in *(B)*.

For an in-depth mode-of-action analysis of CGlcN and to address these potential secondary effects of CGlcN treatment on bacterial cell wall metabolites, we performed targeted metabolomics. To this end, we treated *B. subtilis* at exponential growth phases with a 2-fold MIC (64 μg/mL) of CGlcN for two hours. A detailed description of the experimental procedures can be found in the Supporting Information. In brief, the cells were then harvested by centrifugation and the pellets were resuspended in water and disrupted by sonication. Cell debris was separated from the cytosolic fraction by centrifugation and proteins were removed by acetone precipitation. The acetone was evaporated and samples dissolved in water. The isolated cytosolic fractions were analyzed by LC-MS.

The results of these experiments demonstrate metabolization of CGlcN, thereby yielding the carba-variants of the central cell wall precursor molecule UDP-GlcNAc and the lipidI/II progenitor UDP-MurNAc-5P, namely UDP-CGlcNAc and UDP-CMurNAc-5P (**Fig. 2a,b**). We detected metabolites with a mass difference of 2 Da in the CGlcN treated bacterial cell lysates, reflected by the replacement of the ring oxygen atom with a methylene group (**Fig. 2a,b**). Of note, CGlcN6P was not detectable, indicating a rapid turnover of the compound once taken up through the PTS.

**Figure 2.**
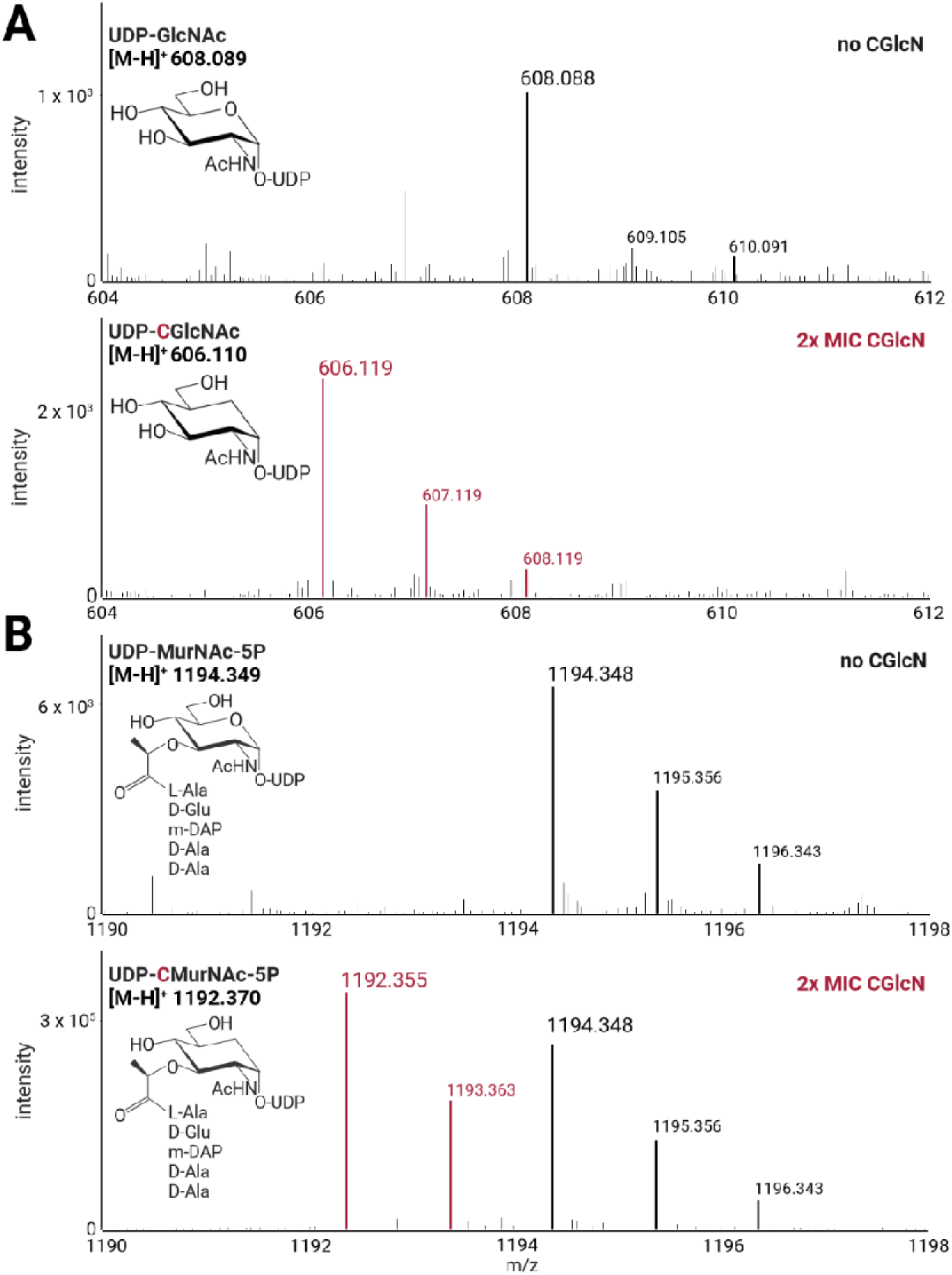
Occurrence of UDP-CGlcNAc and UDP-CMurNAc-5P in a soluble extract of CGlcN-treated *B. subtilis* cells. *(A)* In the presence of CGlcN, UDP-GlcNAc was not identified but a compound with a 2 Da lower mass. Identical MS scans are shown for the control (no CGlcN) and the treated samples (2x MIC CGlcN) at a retention time of 18.7 min. *(B)* In the presence of CGlcN besides UDP-MurNAc-5P a compound with a 2 Da lower mass was detected. MS scans at a retention time of 25.1 min reveal a mixture of UDP-MurNAc-5P and UDP-CMurNAc-5P. Structures and theoretical exact masses are shown. Created with BioRender.com.

We next tested whether the inhibition of certain steps during the peptidoglycan biosynthesis pathway influences the accumulation of specific cell wall precursor molecules, both in the absence and presence of CGlcN. In this regard, we investigated the known antibiotics fosfomycin and vancomycin without or in combination with CGlcN. Fosfomycin inhibits the conversion of UDP-GlcNAc to UDP-MurNAc. Thus, in the presence of fosfomycin UDP-GlcNAc accumulates (**Fig. 3**). When CGlcN was administered together with fosfomycin a strong accumulation of UDP-CGlcNAc was measurable whereas UDP-GlcNAc was no longer detectable. The treatment of *B. subtilis* with vancomycin leads to the accumulation of UDP-MurNAc-5P. In the presence of CGlcN, a considerable amount of UDP-CMurNAc-5P accumulates in addition to UDP-MurNAc-5P (**Fig. 3**). Thus, in contrast to the treatment with fosfomycin, both precursors were detected during vancomycin treatment.

**Figure 3.**
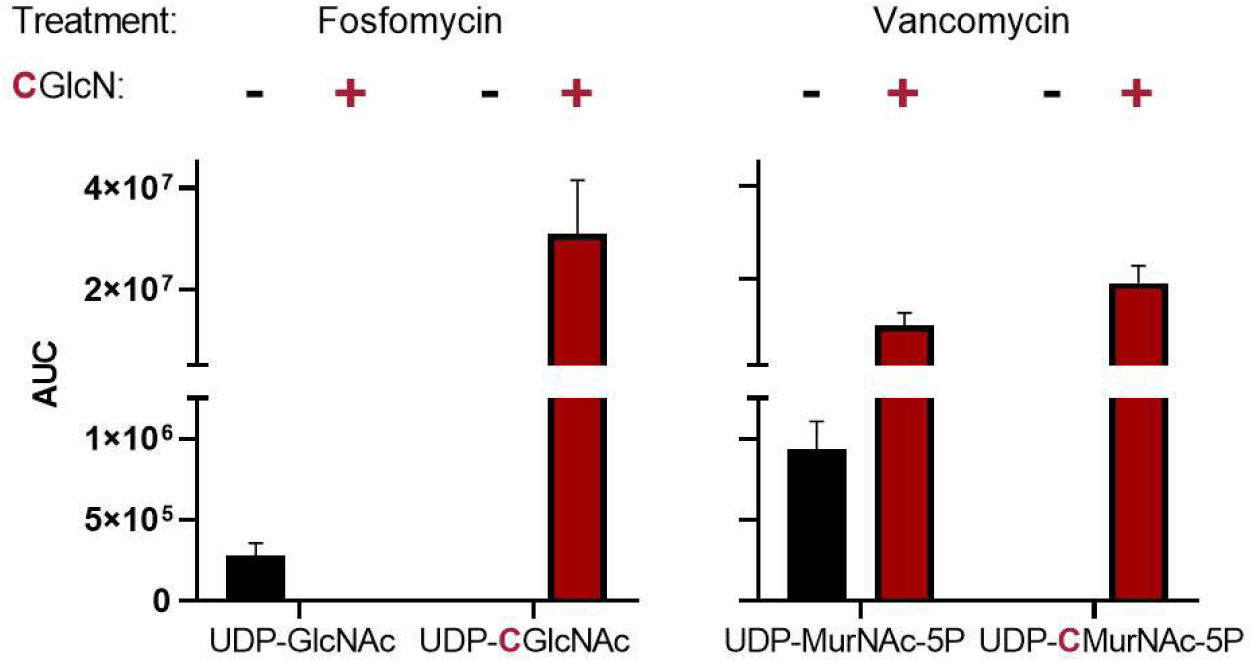
Accumulation of peptidoglycan nucleotide precursors and carbocyclic variants thereof by inhibition with antibiotics. Shown are the relative metabolite concentrations expressed as area under the curve (AUC) values from three independent experiments determined by extracted ion chromatograms with 0.02 Da mass tolerance and baseline adjusted to 500 cps. CGlcN was added at a concentration of 64 μg/ml (2x MIC) and the antibiotic concentrations were 512 μg/ml for fosfomycin and 1 μg/ml vancomycin (2x MIC in all cases as determined for *B. subtilis* grown in CDM medium). Black bars: w/o CGlcN; red bars: w/ CGlcN.

In conclusion, we describe the *in vivo* generation of antimetabolites, generated from CGlcN along the cell wall precursor biosynthetic pathway (**Fig. 4**). These antimetabolites add a second effect to the antibiotic profile of CGlcN, besides its previously shown induction of the *glmS* riboswitch.^3^ Compounds acting on riboswitches are promising antibiotics.^18,19^ However, metabolite analogues of riboswitch activators bear the advantage of secondary effects that add to the initial mode of action.^20^ Further analysis towards carba-variants of lipid I/II are of interest, however this requires developing elaborated protocols to address hydrophobic membrane associated compounds. According to our current model, we believe that the main antibiotic activity relates to the antimetabolite effect of CGlcN and only secondary to *glmS* riboswitch activation. This perception is supported by the characteristics observed from F-CGlcN, which are similar to CGlcN regarding bacterial growth inhibition, PTS dependency and induction of cell envelope stress.^3,10^ In terms of riboswitch activation in vitro they are substantially different though. F-CGlcN also inhibits bacterial growth by metabolization, but whether the extend of antimetabolite generation is identical or differs from the one observed by CGlcN remains elusive. Due to limited availability of the fluoro-compound we could not test it in this study. Taken together, these data demonstrate that CGlcN is not only a putative antibiotic molecule with previously unknown antimetabolite properties, but also a promising tool compound broadly applicable to unravel bacterial cell wall metabolism, e.g., in synergy with other antibiotics or by using carba-variants for targeted labelling of cell wall precursors.

**Figure 4.**
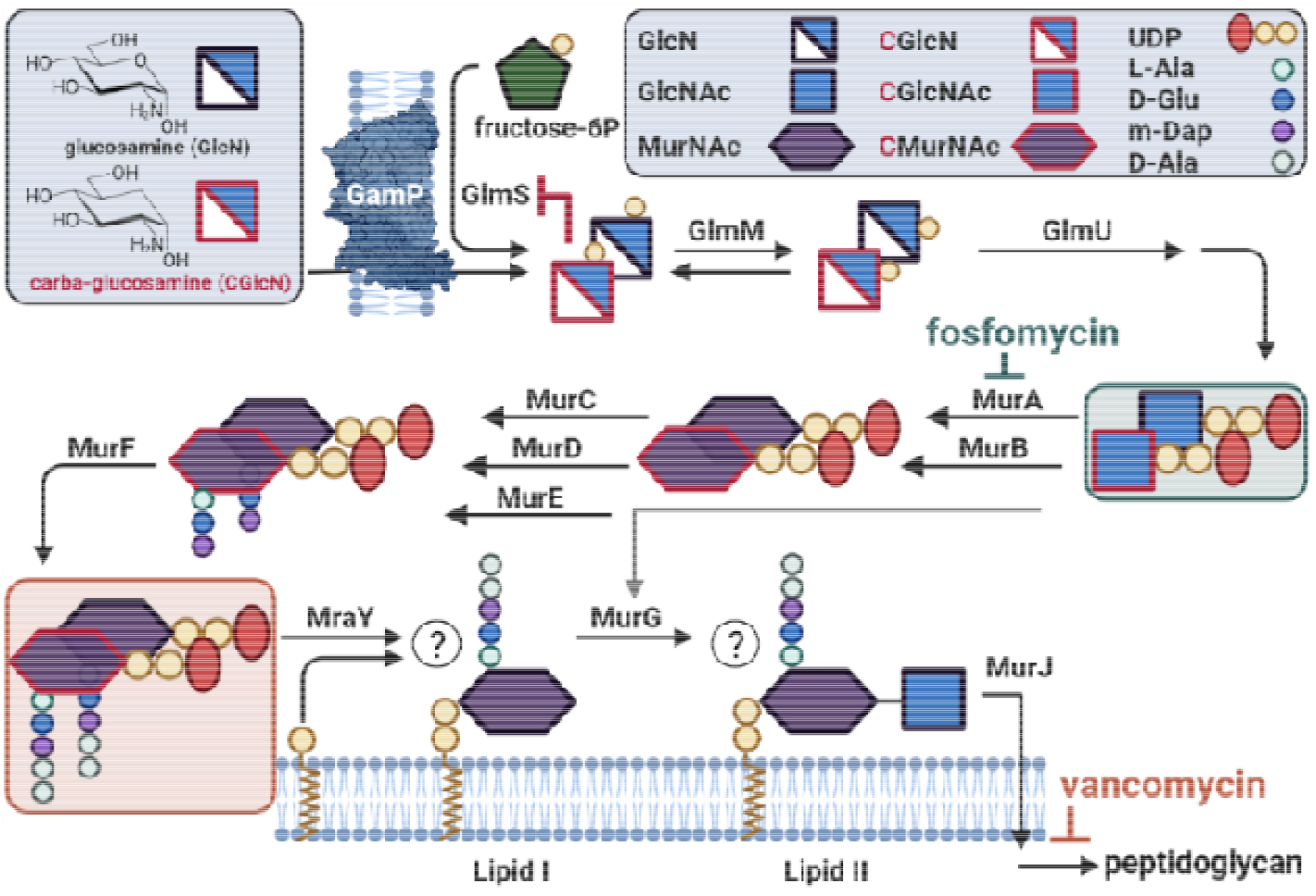
Schematic of the metabolism of glucosamine and CGlcN in *Bacillus subtilis* along the peptidoglycan biosynthetic pathway. The carbocyclic analog is taken up from the growth medium and phosphorylated via the phosphotransferase GamP. The phosphorylated product can interfere with GlmS regulation^2,10^, however also serves as fortuitous substrate of the glucosamine isomerase GlmM and further on in the pathway as substrate of the bifunctional acetyl-/uridylyltransferase GlmU, yielding UDP-CGlcNAc, which subsequently is metabolized via the Mur enzymes up to UDP-CMurNAc-5P. The latter apparently serves as the deadend product. Whether MraY and MurG also accept the carbon derivatives as substrates is currently unclear. Shown are the target steps of the antibiotics fosfomycin and vancomycin used to accumulate the carba-analogs of the peptidoglycan nucleotide precursors. Created with BioRender.com.

## Supporting Information available

Supporting Information contain experimental procedures, materials and methods description and supporting tables.

## Author Contributions

This study was conceived by GM and CM. GM and CM supervised the experiments, MM and SF conducted the metabolomics experiments, evaluated and interpreted results. DaM and AE synthesized the F-CGlcN and examined its biological activity. AB and DiM synthesized CGlcN. HBO supervised the promoter-dependent assays of F-CGlcN. The manuscript was written by GM, CM, MM and SF. All authors have given approval to the final version of the manuscript.

## Acknowledgement

This work was supported by the German Research Foundation (DFG) Project-ID 398967434 — TRR261 to GM, CM, HBO and DiM. The authors also acknowledge Prof G. Bierbaum (University of Bonn) for helpful discussion and support relating to this project. We thank Catherine Schumacher for supporting the promotor assays using F-CGlcN.

## REFERENCES

(1) Sun, Y.; Nitz, M. Syntheses of Carbocyclic Analogues of alpha-D-Glucosamine, alpha-D-Mannose, alpha-D-Mannuronic Acid, beta-L-Idosamine, and beta-L-Gulose. J Org Chem 2012, 77 (17), 7401–7410. https://doi.org/10.1021/jo301240j.

(2) Luense, C. E.; Schmidt, M. S.; Wittmann, V.; Mayer, G. Carba-Sugars Activate the GlmS-Riboswitch of Staphylococcus Aureus. Acs Chem Biol 2011, 6 (7), 675–678. https://doi.org/10.1021/cb200016d.

(3) Schueller, A.; Matzner, D.; Luense, C. E.; Wittmann, V.; Schumacher, C.; Unsleber, S.; Broetz◻Oesterhelt, H.; Mayer, C.; Bierbaum, G.; Mayer, G. Activation of the GlmS Ribozyme Confers Bacterial Growth Inhibition. Chembiochem 2017, 18 (5), 435–440. https://doi.org/10.1002/cbic.201600491.

(4) Babczyk, A.; Wingen, L. M.; Menche, D. Optimized and Scalable Synthesis of Carba◻α◻d◻Glucosamine. Eur. J. Org. Chem. 2020, 2020 (42), 6645–6648. https://doi.org/10.1002/ejoc.202001203.

(5) Staengle, D.; Silkenath, B.; Gehle, P.; Esser, A.; Mayer, G.; Wittmann, V. Carba◻Sugar Analogs of Glucosamine◻6◻Phosphate: New Activators for the GlmS Riboswitch. Chem European J 2023, 29 (3), e202202378. https://doi.org/10.1002/chem.202202378.

(6) Winkler, W. C.; Nahvi, A.; Roth, A.; Collins, J. A.; Breaker, R. R. Control of Gene Expression by a Natural Metabolite-Responsive Ribozyme. Nature 2004, 428 (6980), 281–286. https://doi.org/10.1038/nature02362.

(7) Collins, J. A.; Irnov, I.; Baker, S.; Winkler, W. C. Mechanism of MRNA Destabilization by the GlmS Ribozyme. Gene Dev 2007, 21 (24), 3356–3368. https://doi.org/10.1101/gad.1605307.

(8) Esser, A.; Mayer, G. Characterization of the GlmS Ribozymes from Listeria Monocytogenes and Clostridium Difficile. Chem European J 2023, 29 (3), e202202376. https://doi.org/10.1002/chem.202202376.

(9) Klein, D. J.; Ferré-D’Amaré, A. R. Structural Basis of GlmS Ribozyme Activation by Glucosamine-6-Phosphate. Science 2006, 313 (5794), 1752–1756. https://doi.org/10.1126/science.1129666.

(10) Matzner, D.; Schueller, A.; Seitz, T.; Wittmann, V.; Mayer, G. Fluoro◻Carba◻Sugars Are Glycomimetic Activators of the GlmS Ribozyme. Chem European J 2017, 23 (51), 12604–12612. https://doi.org/10.1002/chem.201702371.

(11) Urban, A.; Eckermann, S.; Fast, B.; Metzger, S.; Gehling, M.; Ziegelbauer, K.; Ru◻bsamen-Waigmann, H.; Freiberg, C. Novel Whole-Cell Antibiotic Biosensors for Compound Discovery. Appl Environ Microb 2007, 73 (20), 6436–6443. https://doi.org/10.1128/aem.00586-07.

(12) Hutter, B.; Fischer, C.; Jacobi, A.; Schaab, C.; Loferer, H. Panel of Bacillus Subtilis Reporter Strains Indicative of Various Modes of Action. Antimicrob Agents Ch 2004, 48 (7), 2588–2594. https://doi.org/10.1128/aac.48.7.2588-2594.2004.

(13) Hutter, B.; Schaab, C.; Albrecht, S.; Borgmann, M.; Brunner, N. A.; Freiberg, C.; Ziegelbauer, K.; Rock, C. O.; Ivanov, I.; Loferer, H. Prediction of Mechanisms of Action of Antibacterial Compounds by Gene Expression Profiling. Antimicrob Agents Ch 2004, 48 (8), 2838–2844. https://doi.org/10.1128/aac.48.8.2838-2844.2004.

(14) Wenzel, M.; Chiriac, A. I.; Otto, A.; Zweytick, D.; May, C.; Schumacher, C.; Gust, R.; Albada, H. B.; Penkova, M.; Krämer, U.; Erdmann, R.; Metzler-Nolte, N.; Straus, S. K.; Bremer, E.; Becher, D.; Brötz-Oesterhelt, H.; Sahl, H.-G.; Bandow, J. E. Small Cationic Antimicrobial Peptides Delocalize Peripheral Membrane Proteins. Proc National Acad Sci 2014, 111 (14), E1409–E1418. https://doi.org/10.1073/pnas.1319900111.

(15) Gaugué, I.; Oberto, J.; Plumbridge, J. Binding of NagR and GamR to Their DNA Targets. Mol Microbiol 2014, 92 (1), 100–115. https://doi.org/10.1111/mmi.12544.

(16) Gaugué, I.; Oberto, J.; Putzer, H.; Plumbridge, J. The Use of Amino Sugars by Bacillus Subtilis: Presence of a Unique Operon for the Catabolism of Glucosamine. Plos One 2013, 8 (5), e63025. https://doi.org/10.1371/journal.pone.0063025.

(17) Deutscher, J.; Francke, C.; Postma, P. W. How Phosphotransferase System-Related Protein Phosphorylation Regulates Carbohydrate Metabolism in Bacteria. Microbiol Mol Biol R 2006, 70 (4), 939–1031. https://doi.org/10.1128/mmbr.00024-06.

(18) Mulhbacher, J.; Brouillette, E.; Allard, M.; Fortier, L.-C.; Malouin, F.; Lafontaine, D. A. Novel Riboswitch Ligand Analogs as Selective Inhibitors of Guanine-Related Metabolic Pathways. Plos Pathog 2010, 6 (4), e1000865. https://doi.org/10.1371/journal.ppat.1000865.

(19) Howe, J. A.; Wang, H.; Fischmann, T. O.; Balibar, C. J.; Xiao, L.; Galgoci, A. M.; Malinverni, J. C.; Mayhood, T.; Villafania, A.; Nahvi, A.; Murgolo, N.; Barbieri, C. M.; Mann, P. A.; Carr, D.; Xia, E.; Zuck, P.; Riley, D.; Painter, R. E.; Walker, S. S.; Sherborne, B.; Jesus, R. de; Pan, W.; Plotkin, M. A.; Wu, J.; Rindgen, D.; Cummings, J.; Garlisi, C. G.; Zhang, R.; Sheth, P. R.; Gill, C. J.; Tang, H.; Roemer, T. Selective Small-Molecule Inhibition of an RNA Structural Element. Nature 2015, 526 (7575), 672–677. https://doi.org/10.1038/nature15542.

(20) Krajewski, S. S.; Isoz, I.; Johansson, J. Antibacterial and Antivirulence Effect of 6-N-Hydroxylaminopurine in Listeria Monocytogenes. Nucleic Acids Res 2017, 45 (4), 1914–1924. https://doi.org/10.1093/nar/gkw1308.

