## Supporting Information for "Metabolization of α-D-carba-glucosamine in vivo generates antimetabolites of cell wall precursors"

**Experimental procedures, materials and methods**

**Carba-sugars**

CGlcN and F-CGlcN were synthesized as published previously^1,2^.

**MIC determination**

Lysogenic broth (LB) medium (5 ml) was inoculated with *B. subtilis* 168 wildtype and the culture was grown until OD_600_ of ~1. Bacteria were diluted 1:100 in 5 ml in chemically defined medium^3^ and incubated overnight at 37°C and 130 rpm. A two-fold serial dilution of F-CGlcN, fosfomycin or vancomycin in CDM was prepared in a U-shaped 96-well plate including a growth control (bacterial suspension without F-CGlcN or antibiotic). The plate was then inoculated with 1x10^6 CFU/ml for F-CGlcN or 1x10^5 CFU/ml for fosfomycin and vancomycin and incubated at 37 °C for 16-20 hours or 24 hours, respectively. Subsequently the MIC was determined.

**Growth curves**

*B. subtilis* 168 wildtype was pre-cultured overnight in LB-medium at 37 °C and 130 rpm. The next day, 5 ml CDM were inoculated 1:100 with bacterial cells and grown till OD_600_ of 0.5-0.7. The culture has been diluted to OD_600_=0.1 and cells were transferred into U-shaped 96-well plates. Growth was monitored in the presence of F-CGlcN (0-300 µM). The plate was incubated for 10 hours at 37°C while shaking for 3 sec prior to each measurement. The OD_600_ was measured every 3 min using an EnSpire plate reader (PerkinElmer).

**ΔPTS-dependent growth curves**

*B. subtilis* 168 wildtype and mutant strains were obtained from *B. subtilis* genetic stock center (Table 1). They were incubated in CDM–erythromycin (3 ml) at 37°C overnight. The precultures were diluted to OD_600_=0.1 and F-CGlcN was add to a final concentration of 300 µM (2xMIC). Growth curves were monitored as described above using a Sunrise plate reader (Tecan).

**Cell stress promotor assay**

*B. subtilis* reporter strains (Table 2) were pre-cultured overnight in LB–erythromycin (10 mL) at 37 °C. Subsequently the culture was diluted to OD_600_=0.05 in CDM (5 mL) and grown until OD_600_≈0.2 was reached. A serial dilution of CGlcN was inoculated to a final OD_600_ of 0.01. The cells were incubated for 4 h at 37°C. Subsequently citrate buffer (60 µL, 0.1 M, pH 5) containing luciferin (2 mM, Serva, Heidelberg, Germany) was added and luminescence was measured using a Sunrise plate reader (Tecan).

**Cultivation conditions for metabolomic studies**

*B. subtilis* 168 wildtype was grown in CDM without glucose at 37°C under continuous shaking (170 rpm). The main culture was inoculated with an overnight culture to an OD_600_ of 0.1. Bacterial cells were grown to exponential growth phase (OD_600_ of 0.3-0.6). The culture was split into 50 ml aliquots and these were treated differentially with CGlcN 64 µg/ml (2x MIC) alone or in combination with 2x MIC of either of the antibiotics fosfomycin (512 µg/ml) and vancomycin (1 µg/ml). MICs of fosfomycin and vancomycin in CDM were determined as described above. As control, bacteria were grown without antibiotics or being treated with only one of the antibiotics. Cells were harvested by centrifugation (10 min, 3500 x g, 4°C) two hours after the addition of the carba-sugar and the indicated antibiotic. OD_600_ bevor harvest was measured.

**LC-MS sample preparation**

Pellets were thawed on ice and resuspended in 2 ml H_2_O by vortexing. Cells were then disrupted by sonication. Cell debris was separated from soluble fraction by centrifugation at 15000 x g for 30 min. The supernatant was transferred into a new tube. Proteins were precipitated by adding 800 µl ice-cold acetone to 200 µl supernatant. The samples were incubated for 30 min at -20°C. Then samples were centrifuged for 10 min at 11000 x g. After centrifugation the supernatant containing the metabolites was transferred into a new tube. Acetone evaporated overnight under the fume hood. Samples were freeze-dried and stored at -80°C until analysis. The dried samples were dissolved in H_2_O_dd_. In regard of the measured OD_600_ the samples were adjusted with the corresponding amount of H_2_O_dd_.

**LC-MS analysis und data evaluation**

Analysis was performed with an electro-spray ionization-time-of-flight (ESI-TOF) mass spectrometer (MicrOTOF II; Bruker Daltonics) operated in positive ion-mode (200-1600 m/z) that was connected to an UltiMate 3000 high performance liquid chromatography (HPLC) system (Dionex). A Gemini C18 column (150 x 4.6 mm, 5 μm, 110 Å, Phenomenex) was used and a HPLC elution program with 0.2 ml/min flow rate, 5-minute wash step with 100% buffer A (0.1% formic acid with acid with 0.05% ammonium formate), continued by a linear gradient over 30 min to 40% eluent buffer B (buffer B = 100% acetonitrile), 5-min time delay of the gradient and a 5 min re-equilibration. 5 µL of each sample was injected.

The evaluation was performed using extracted ion chromatograms (accuracy 0.02 Da), the AUC (baseline 500 cps) was determined using the OpenMS^4^ based tool for metabolomics data analysis UmetaFlow^5^ with its graphical user interface (<https://github.com/axelwalter/umetaflow-gui>). The AUCs obtained from UmetaFlow were presented in Prism 8 (GraphPad).

The determination of the AUC was not only carried out with the [M+H]^+^ mass but also with the adducts [M+Na]^+^, relevant in the case of UDP-(C)-GlcNAc and the doubly charged mass [M+H]^2+^, relevant in the case of UDP-(C)-MurNAc-5P. These adducts and charges make up a significant part of the corresponding metabolites. In addition, the relevance of the AUC was ensured by specifying the retention time. An overview of the metabolites is shown in table 3.

**Supporting Tables:**

**Table 1. List of *B. subtilis* knock-out mutant strains**

| *B. subtilis* strain | *B. subtilis* genetic stock center catalogue number |
| --- | --- |
| ΔptsH::ermR | BKE13900 |
| ΔgamP::ermR | BKE02350 |
| ΔnagP::ermR | BKE07700 |

**Table 2. List of stress inducible *B. subtilis* strains**

| *B. subtilis* strain | Stress-inducible promotor |
| --- | --- |
| *B. subtilis* 1S34 | none |
| *B. subtilis* 1S34 pS 63 | *helD*, synonym *yvgS* (inhibition of RNA synthesis) |
| *B. subtilis* 1S34 pS 72 | *bmrC*, synonym *yheI* (inhibition of protein synthesis) |
| *B. subtilis* 1S34 pS 77 | *yorB* (inhibition of DNA synthesis or DNA damage) |
| *B. subtilis* 1S34 pS 107 | *ypuA* (cell envelope stress) |

**Table 3. Relevant masses for EIC evaluation of LC-MS samples**

| Metabolite | neutral monoisotopic mass | [M+H]^+^ | [M+Na]^+^ | [M+H]^2+^ | Retention time [min] |
| --- | --- | --- | --- | --- | --- |
| UDP-GlcNAc | 607.0816 | 608.0889 | 630.0708 | 304.5481 | 16.0-22.0 |
| UDP-Carba-GlcNAc | 605.1023 | 606.1096 | 628.0915 | 303.5584 | 16.0-22.0 |
| UDP-MurNAc-5P | 1193.3414 | 1194.3487 | 1216.3307 | 597.6780 | 24.5-27.0 |
| UDP-Carba-MurNAc-5P | 1191.3622 | 1192.3694 | 1214.3514 | 596.6884 | 24.5-27.0 |

**References**

1. Babczyk, A.; Wingen, L. M.; Menche, D. Optimized and scalable synthesis of carba-α-D-glucosamine. European Journal of Organic Chemistry 2020, 2020 (42), 6645-6648. DOI: https://doi.org/10.1002/ejoc.202001203

2. Matzner, D.; Schuller, A.; Seitz, T.; Wittmann, V.; Mayer, G. Fluoro-carba-sugars are glycomimetic activators of the glmS ribozyme. Chemistry 2017, 23 (51), 12604-12612. DOI: 10.1002/chem.201702371 From NLM Medline.

3. Van de Rijn I., Kessler R. E. Growth Characteristics of Group A Streptococci in a New Chemically Defined Medium. Infect Immun 27:444–448. (1980).

4. Rost HL, Sachsenberg T, Aiche S, Bielow C et al. OpenMS: a flexible open-source software platform for mass spectrometry data analysis. Nat Meth. 2016; 13, 9: 741-748. doi:10.1038/nmeth.3959.

5. Kontou EE, Walter A, Alka O, Pfeuffer J, Sachsenberg T, Mohite O, et al. UmetaFlow: An untargeted metabolomics workflow for high-throughput data processing and analysis. ChemRxiv. Cambridge: Cambridge Open Engage; 2022; This content is a preprint and has not been peer-reviewed. [10.26434/chemrxiv-2022-z0t4g-v2](https://doi.org/10.26434/chemrxiv-2022-z0t4g-v2).
